# The marine natural microbiome mediates physiological outcomes in host nematodes

**DOI:** 10.1101/2023.05.10.540299

**Authors:** Yiming Xue, Yusu Xie, Xuwen Cao, Liusuo Zhang

## Abstract

Nematodes are the most abundant metazoans in marine sediments, many of which are bacterivores, however how habitat bacteria effects physiological outcomes in marine nematodes remains largely unknown. Here, we used a *Litoditis marina* inbred line to assess how native bacteria modulates host nematode physiology. We characterized seasonal dynamic bacterial compositions in *L. marina* habitats, and examined the impacts of 448 habitat bacteria isolates on *L. marina* development, then focused on HQbiome with 73 native bacteria, of which we generated 72 whole genomes sequences. Unexpectedly, we found that the effects of marine native bacteria on the development of *L. marina* and its terrestrial relative *Caenorhabditis elegans* were significantly positively correlated. Next, we reconstructed bacterial metabolic networks and identified several bacterial metabolic pathways positively correlated with *L. marina* development (e.g., ubiquinol and heme *b* biosynthesis), while pyridoxal 5’-phosphate biosynthesis pathway was negatively associated. Through single metabolite supplementation, we verified CoQ_10_, heme *b*, Acetyl-CoA, and acetaldehyde promoted *L. marina* development, while vitamin B6 attenuated growth. Notably, we found that only four development correlated metabolic pathways were shared between *L. marina* and *C. elegans*. Furthermore, we identified two bacterial metabolic pathways correlated with *L. marina* lifespan, while a distinct one in *C. elegans*. Strikingly, we found that glycerol supplementation significantly extended *L. marina* but not *C. elegans* longevity. Moreover, we comparatively demonstrated the distinct gut microbiota characteristics and their effects on *L. marina* and *C. elegans* physiology. Our integrative approach will provide a microbe–nematodes framework for microbiome mediated effects on host animal fitness.

## Introduction

Multicellular organisms, including animals and plants, interact closely with diverse and abundant microbial communities in their natural habitats^1–3^. Both the microbes physically associated with the host and the animal gut microbiota influence host physiology and fitness^2–4^. The microbiota communities modulate essential biological processes such as growth, development, reproduction, longevity, and responses to biotic and abiotic stresses, in their host animals and plants^5–7^. Changes in environmental bacteria can affect the behaviors of zebrafish such as locomotion^8^ and anxiety-related behavior^9^. Experimental studies in mouse models have demonstrated substantial impacts of the gut microbiota on neuron development, angiogenesis, immunity, and disease^10,11^. The dysbiosis of gut microbiota has been associated with a wide spectrum of human diseases such as obesity, diabetes, cancer, and neurological disorders^12^.

Among the model organisms utilized to study the interactions between microbiota and host animals, the terrestrial nematode *Caenorhabditis elegans* is an excellent model to study microbiome-mediated functions, due to its unique characteristics such as a short generation time, clear genetic background, and advanced functional genomics resources^13^. *C. elegans* encounters different bacteria in its native habitats, which could be potentially used as dietary resources, some of which colonize the worm’s gut^14^. It was reported that nearly 80% of the >550 habitat bacterial isolates could individually support *C. elegans* development^15^. The effects of several microbial metabolites on host physiology have been investigated in *C. elegans*, e.g., vitamins^16–18^, siderophores^19,20^, and neurotransmitters^21^. Genome-scale metabolic models were recently reconstructed for 77 members of *C. elegans* natural habitat bacteria, revealing key bacterial metabolic traits related to gut bacterial colonization and host fitness^22^. Recently, the gut microbiota of *C. elegans* was reported to be shaped by both host genetics and environmental factors^14,22–24^.

Although both nematodes and bacteria are dominant organisms in marine sedimentary ecosystems, and many marine nematodes are bacterivores, however how native bacteria affect the essential biological processes of marine nematodes is largely unexplored. *Litoditis marina*, a marine free-living bacterivore nematode, is widely distributed and plays an important role in marine ecosystems^25–27^. We have recently established *L. marina* as a promising marine model organism, including a short generation time, easy to be cultured in the lab, multiple inbred lines with clear genetic background, high-quality genome assembly, annotations, and functional genomics resources^26,27^. It was reported that the gut microbiome of cryptic species of *L. marina* is highly diverse and species-specific^25^. Native bacteria might provide nutrient resources to *L. marina*, and the natural microbiome is affected by global climate change, however, the composition of the *L. marina* habitat microbial communities and the natural microbiome-mediated functions on nematode fitness are largely unknown.

In this study, we characterized the native bacterial community compositions and isolated 539 bacterial strains from *L. marina* natural habitats. Next, the impacts of 448 single native bacterial isolates on nematode development were assessed, and a representative set (73 bacteria, termed HQbiome) was further examined for their effects on host physiology, of which 72 whole genome sequences were generated. Unexpectedly, we found that the effects of marine native bacteria on the development of *L. marina* and its terrestrial relative *C. elegans* were significantly positively correlated. Based on the whole genome sequences of HQbiome, we reconstructed bacterial metabolic networks and assessed their metabolic capacities. Based on HQbiome bacterial metabolic networks reconstruction with random forest regression analysis, together with single metabolite supplementation, we demonstrated that CoQ_10_, heme *b*, acetyl-CoA, and acetaldehyde promoted *L. marina* development, while vitamin B6 attenuated growth. Notably, we found that only four growth development correlated metabolic pathways were shared between *L. marina* and *C. elegans*. Strikingly, we found that glycerol supplementation significantly extended *L. marina* but not *C. elegans* longevity. Moreover, we demonstrated the gut microbiota characteristics and bacterial effects on the physiology of both *L. marina* and *C. elegans*.

## Results

### The natural bacterial microenvironment of *L. marina*

To characterize the bacterial community composition that *L. marina* encounters in its natural environment, we performed full-length 16S rRNA gene amplicon sequencing of 80 intertidal sediment samples from the *L. marina* habitats (Huiquan Bay, Qingdao, China) over a thirteen-month period (Fig. S1). A total of 5,783 operational taxonomic units (OTUs) were identified (Table S1A). Of these, the dominant phyla were Proteobacteria (46.33%), Bacteroidetes (22.03%), Cyanobacteria (11.11%), Actinobacteria (6.30%), Verrucomicrobia (3.13%), Acidobacteria (2.96%), Planctomycetes (1.78%), and Gemmatimonadetes (1.60%) (Fig. 1a and Fig. S2; Table S1B,C). Notably, Proteobacteria and Bacteroidetes are also the most abundant phylum under *C. elegans* natural habitats^15^. As the most abundant phylum, Proteobacteria were mostly enriched with classes such as Gammaproteobacteria (28.18%), Deltaproteobacteria (10.91%), Alphaproteobacteria (5.93%), and Betaproteobacteria (0.87%). We found that phylum Cyanobacteria was significantly abundant in winter (Fig. 1a), which was previously reported to be well adapted to cold environments^28,29^. Of the 1821 identified genera, the most abundant genus was *Woeseia*, presenting in all 80 samples, and being the only genus with more than 10% mean abundance (11.80%, Fig. 1b; Table S1D). Moreover, assessments of α-diversity revealed a significantly lower richness and diversity of the natural microbiome in the coldest month (January) compared with that of other seasons (Fig. S3; *P_FDR_* < 0.05, Kruskal‒Wallis test, Wilcoxon test). Abundance–occupancy analyses identified six OTUs that were persistent in all samples (Fig. S4; Table S1E). Only 47 (0.98%) OTUs were found in ≥ 70 samples, but these were the dominant bacteria and accounted for 44.32% of the total reads sequenced (Table S1E), being dominated by Gammaproteobacteria (39.39%), Bacteroidetes (20.91%), Cyanobacteria (17.16%), Deltaproteobacteria (12.89%), Actinobacteria (5.73%), and Gemmatimonadetes (1.59%). Their ubiquitous presence and high abundance indicated that these taxa were likely core members of the environmental bacteria of *L. marina*. About 85.60% of all OTUs were found in < 10 samples (4950/5783 OTUs, Table S1E), of which 50.79% OTUs (2937/5783 OTUs) were unique to one sample. Meanwhile, there was large seasonality in the less abundant bacteria (Fig. 1a and Fig. S3). The abundance–occupancy relationships of each phylum were shown in Fig. S4.

**Fig. 1.**
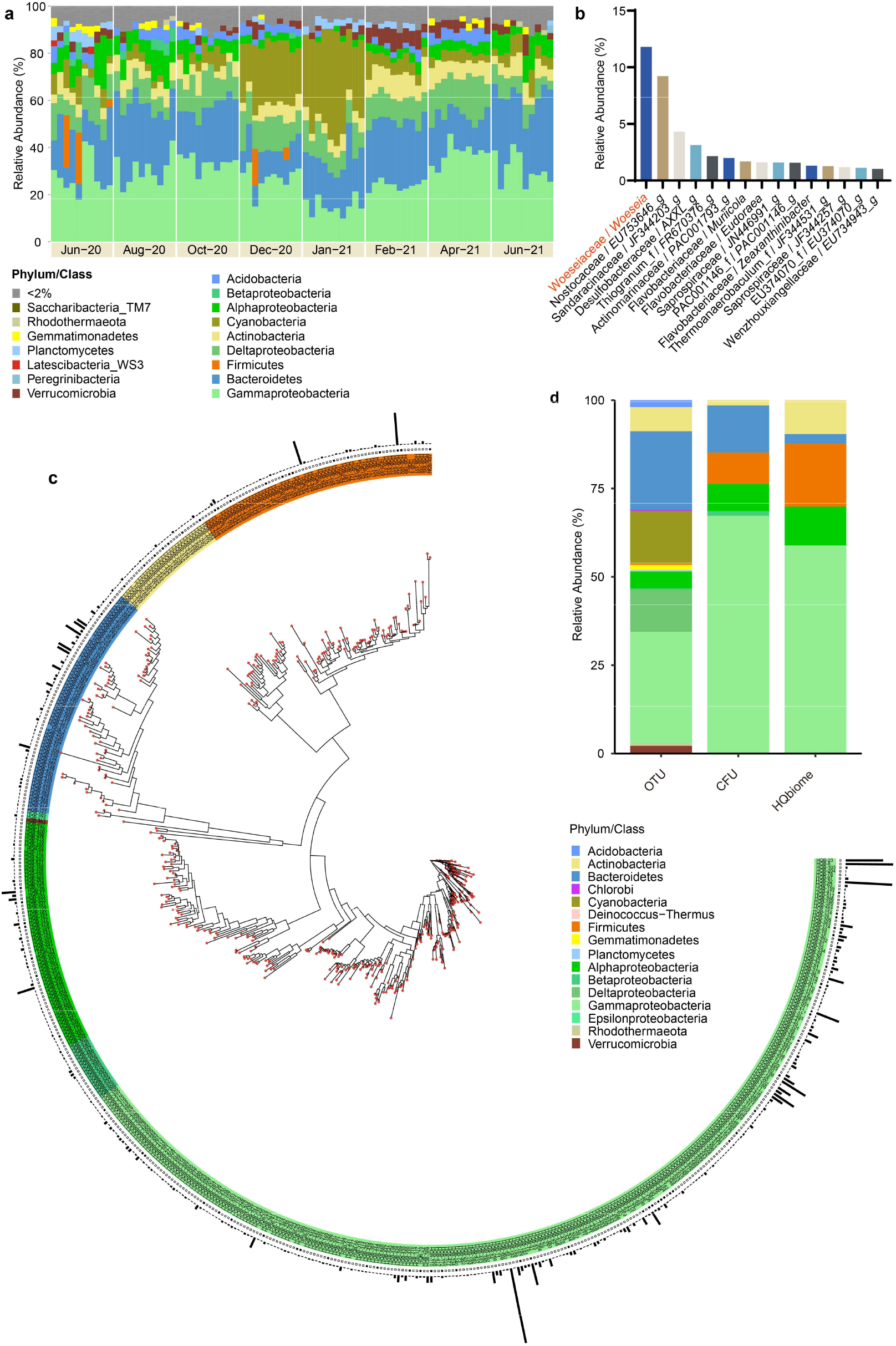
The bacterial composition in *L. marina* natural habitat. **a,** Phylum-level distribution of the *L. marina* natural habitat microbiome through cultivation-independent full-length 16S amplicon sequencing. Proteobacteria are shown at the class level. Relative abundance (RA) less than 2% are grouped into “< 2%”. **b,** Abundant bacterial genera in *L. marina* habitats (greater than 1% mean abundance are shown). **c,** The taxonomic diversity of 539 cultivation-dependent isolates. From inner to outer, the data mapped onto the tree is HQbiome strains (black squares) and the relative abundance of CFUs. Taxonomic assignment and phylogenetic tree inference were based on full-length 16S rRNA gene sequences. **d**, Phylum-level distribution of habitat microbiome (OTUs > 0.1% RA), culture collection (abundance of 3822 CFUs), and HQbiome (73 isolates).

### Impact of natural bacteria on the development of *L. marina*

To ask how individual habitat bacteria may affect the development and longevity of *L. marina*, we isolated bacteria on simple marine bacterial growth media from sediment samples where *L. marina* was collected. A total of 3,822 colony-forming units (CFUs) were obtained and taxonomically characterized by sequencing the bacterial 16S rRNA gene (Table S2A), resulting in a total of 539 unique CFUs (Table S2B), belonging to Gammaproteobacteria (67.06%), Alphaproteobacteria (7.74%), Betaproteobacteria (1.28%), Bacteroidetes (13.34%), Firmicutes (8.84%), and Actinobacteria (1.49%) (Fig. 1c,d). These bacteria isolates were categorized into 10 classes, 34 orders, 68 families, and 192 genera (Table S2B), among which, all phyla, seven classes, 24 orders, 45 families, 85 genera, and 51 species overlapped with our cultivation-independent full-length 16S rRNA community profiling, respectively (Table S2C). The majority of 539 CFUs were members of the genera *Vibrio*, *Photobacterium*, *Pseudoalteromonas*, *Shewanella*, *Olleya*, *Bacillus*, and *Polaribacter*, most of which were also reported from other marine bacterial cultures, which might represent the dominant cultivable bacteria genera in marine and coastal ecosystems^30^. Notably, our cultivation-based collection expanded the diversity of the existing collection, and 28 of them were candidates for new species which had a 16S rRNA gene sequence with ≤ 97% identity to their closest references (Table S2B).

To ask how natural bacteria affect marine nematode development, we fed *L. marina* with individual bacterial isolates (448 of 539 isolates examined in this project). By applying *k*-means clustering, 121 bacterial strains were categorized as generally “beneficial” (fast growth, percentage of stage 4 larvae (L4) in day 5: ≥ 45.71%), 113 were categorized as “intermediate” (moderate growth, percentage of L4 in day 5: 18.57% ∼ 45.71%), and 214 were “detrimental” (slow growth or active killing, percentage of L4 in day 5: < 18.57%) (Fig. 2, Fig. S5 and Fig. S6; Table S2D). Nearly twice as many strains were classified as “detrimental” (47.66%) compared to “beneficial” (27.17%) (Fig. 2a). Specific taxa tend to have predominantly beneficial or detrimental impacts on *L. marina*, for instance, most Alphaproteobacteria (e.g., Rhodobacteraceae) and Betaproteobacteria (e.g., Comamonadaceae) strains were beneficial to *L. marina*, while Actinobacteria (e.g., Micrococcaceae and Nocardiaceae), Firmicutes (e.g., Planococcaceae and Bacillaceae), Bacteroidetes (e.g., Flavobacteriaceae), and several Gammaproteobacteria (e.g., Moraxellaceae, Psychromonadaceae, and Vibrionaceae) showed detrimental impacts (Fig. S7). Note that many of these “detrimental” taxa such as Microbacteriaceae and Colwelliaceae are pathogens of *C. elegans* (see Supplementary Results S1) and other animals^15^. We found a significant phylogenetic signal for the effects on nematode growth (Pagel’s *λ* = 0.411, *P* = 3×10^−1^^3^, Abouheif’s *C*_mean_ = 0.192, *P* = 0.001. The observed phylogenetic signal was robust also to subsampling. Fig. S8; Table S2E). However, the distribution of the impacts on *L. marina* development within the certain phylogenetic tree of our culture collection revealed discrepancies between closely related strains, i.e., different strains belonging to the same genus exhibited variant effects, for example, members in Vibrionaceae, Shewanellaceae, Rhodobacteraceae, and Flavobacteriaceae (Fig. S7).

**Fig. 2.**
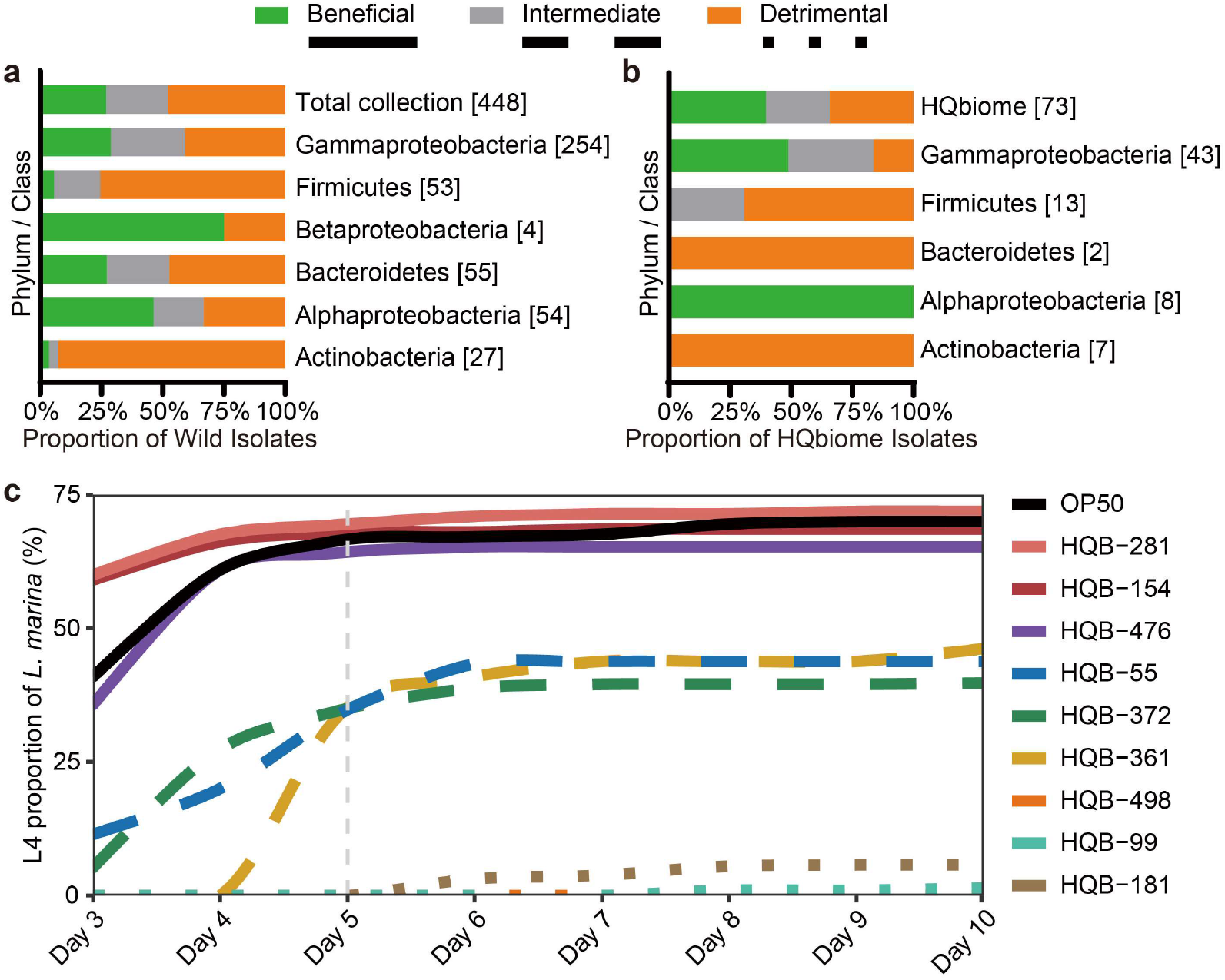
Impact of natural bacterial isolates on *L. marina* development. **a**, Category characterization of the 448 cultured bacteria isolates. **b,** Category characterization of the 73 HQbiome bacteria strains. The clustering of the impacts of natural bacterial isolates on *L. marina* growth rate was analyzed by *k*-means clustering of the proportion of stage 4 larvae (L4) of *L. marina* after 5 days post-hatching (*k* = 3, n = 1,000). Isolate numbers are shown in brackets. **c**, Percentage of L4 of *L. marina* fed with HQbiome bacteria isolates. Beneficial, intermediate, and detrimental bacteria were highlighted in solid, dashed, and dotted lines, respectively. *E. coli* OP50 was shown as a control.

We next focused on 72 taxonomically and functionally representative of the above-mentioned 448 isolates, together with *W. oceani*, which was not isolated by our cultivation-based screening, while is highly represented by our cultivation-independent survey (Fig. 1b), these 73 bacteria were termed HQbiome (Fig. 2b,c; Supplementary Results S2; Table S2F and Table S5). Of the 73 strains in HQbiome, 29 (39.73%) were “beneficial” to *L. marina*, including Alphaproteobacteria and Gammaproteobacteria, while 25 strains (34.25%) belong to the “detrimental” category, most of which were Actinobacteria, Bacteroidetes, and Firmicutes (Fig. 2b). The remaining 19 strains were “intermediate”, belonging to Gammaproteobacteria and Firmicutes. Notably, over 61% of *L. marina* fed with Alphaproteobacteria reached L4 on day 5, whereas Actinobacteria and Bacteroidetes strains significantly attenuated animal growth (Fig. 2b).

### Whole genome sequencing and genome-scale metabolic networks reconstruction and validation

We generated whole genome sequences (WGS) for 72 bacterial isolates from HQbiome. Of which, 15 were sequenced using both Oxford Nanopore Technologies (ONT) and Illumina technologies, one with both Pacific Biosciences (PacBio) and Illumina technologies, resulting in either a single-circular chromosome (9 strains) or a single chromosome with 1 to 6 plasmids (Table S2F). The remaining 56 strains were sequenced with Illumina only, resulting in assemblies with 8 to 211 contigs per genome.

For five genera and two families (*Vibrio*, *Shewanella*, *Pseudoalteromonas*, *Pseudomonas*, *Psychrobacter*, Bacillaceae, and Rhodobacteraceae), each including more than three isolates, we identified substantial inter-specific genome variation (Fig. S9). Based on HQbiome WGS, we constructed a phylogenetic tree and identified unidentical consensus phylogeny compared to the phylogenetic tree inferred from the full-length 16S rRNA gene (Fig. 3a and Fig. S10). Especially, isolates belonging to Bacillaceae clustered well in the WGS phylogenetic tree.

**Fig. 3.**
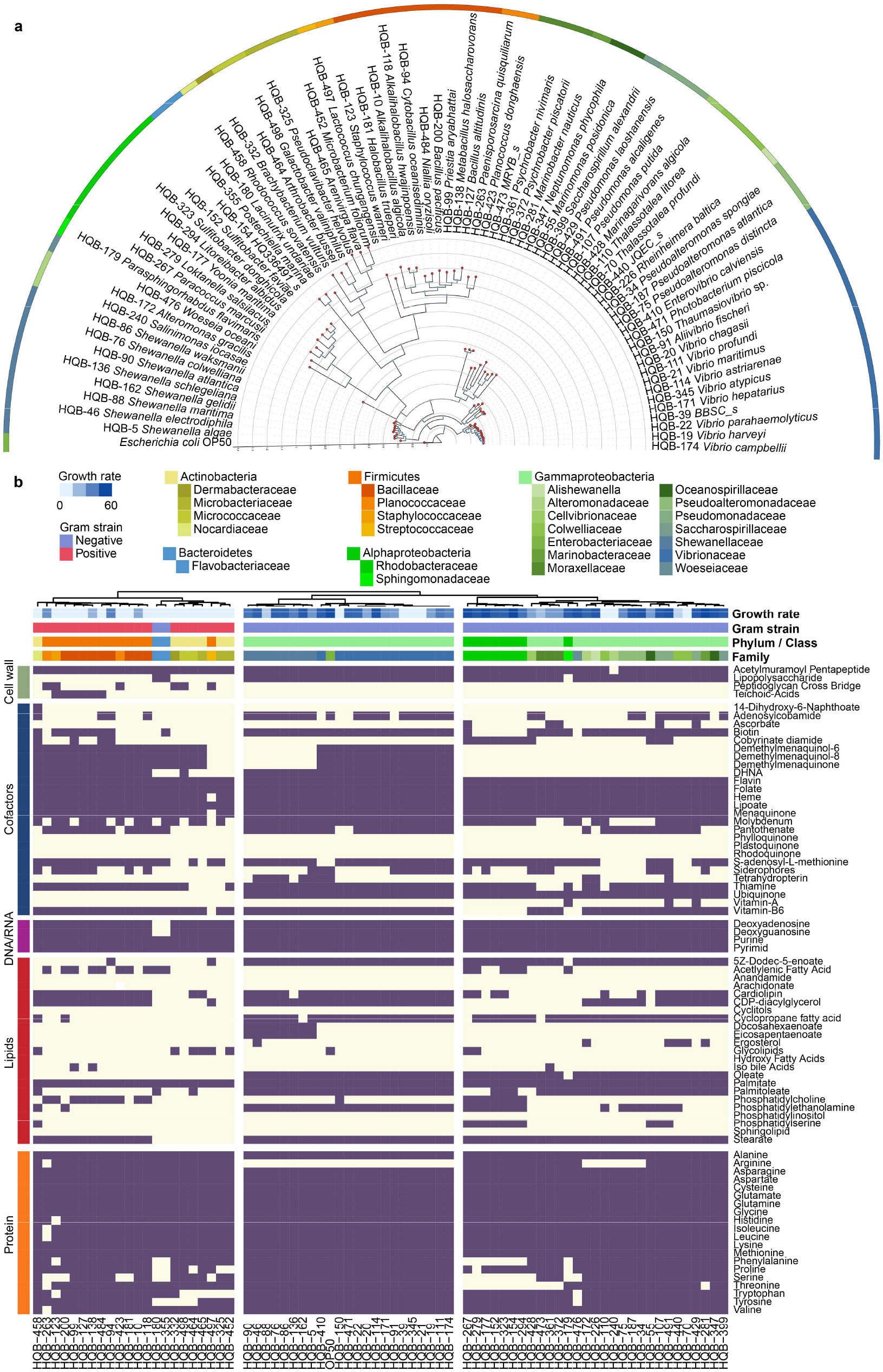
Phylogenetics and the metabolic capacity of HQbiome bacteria isolates. **a,** Phylogenetic tree of HQbiome bacteria isolates based on whole genome sequences. The outer ring represents Family level taxonomy of isolates. **b**, Metabolic competences of the HQBiome strains. Multiple pathways corresponding to the same function were grouped according to the MetaCyc pathway database (Table S3C) and a pathway was considered positive (purple) if one of the associated pathways was confirmed by gapseq. The clustering distance and method were “Euclidean” and “ward.D2”, respectively. *L. marina* growth rate (percentage of L4 after 5 days post-hatching), organism Gram-stain and phylum/class/family are indicated by several annotation columns at the top of the matrix.

To explore metabolic capacities within HQbiome, we reconstructed genome-scale metabolic models (GEMs) using the gapseq pipeline (see Supplementary Results S3; Table S3A). To validate the GEMs, we experimentally quantified the ability of *Escherichia coli* OP50 and four *vibrio* strains to utilize 71 carbon sources using the BIOLOG assay, resulting in 73.20% overlap with the curated models (Fig. S11, Table S3N), which was similar to the previous study^22^. In addition, we evaluated the quality of the GEMs by testing if the models can recapitulate carbon source utilization as indicated in the ProTraits database and found that the correct prediction rate is about 90% (37/41), highlighting the high quality of our reconstructed metabolic models (Fig. S12; Table S3N). Moreover, quality assessment tests with all GEMs were performed with MEMOTE^31^, yielding overall high consistencies: on average, 100% stoichiometric consistency, 99.92% reactions mass-balanced, 99.97% reactions charge-balanced, and 100% metabolite connectivity.

### Metabolic capacity of HQbiome strains

Hierarchical clustering of the genome-informed metabolic potential reflected extensive variability across the HQbiome isolates (Fig. 3b; Table S3C; see Supplementary Results S4 for more details), exhibiting three main clusters: one with generally “detrimental” effects on *L. marina* development, comprising Actinobacteria, Bacteroidetes, and Firmicutes (left), and two with generally “beneficial” effects on *L. marina* growth, including Alphaproteobacteria and Gammaproteobacteria (middle and right). In general, we observed that the basic metabolites essential for their host animals, including nucleotides, essential amino acids, and most cofactors, were found among the “beneficial” isolates (Fig. 3b). Strikingly, the cluster with detrimental effects on *L. marina* development lacked the metabolic potential of several lipids (e.g., oleate, stearate, palmitoleate, cyclopropane fatty acid, and 5Z-dodec-5-enoate) and cofactors (e.g., ubiquinone and biotin). The attenuated *L. marina* development induced by isolates from the “detrimental” cluster could be due to the lack of the above-mentioned metabolic capability for lipids and cofactors synthesis.

### Bacterial metabolites modulate *L. marina* development and longevity

Focusing on HQbiome-1 (“beneficial” to *L. marina*: 29 isolates from HQbiome) and HQbiome-2 (“detrimental” to *L. marina*: 25 isolates from HQbiome) (also see Supplementary Results S5), we identified bacterial metabolic pathways correlated with *L. marina* development through random forest regression analysis (Fig. 4a; Table S3E). Especially, several bacterial metabolic pathways were significantly positively correlated with *L. marina* development, including ubiquinol (Coenzyme Q (CoQ): CoQ_8_, CoQ_9_, CoQ_7_, and CoQ_10_ (PWY-6708, PWY-5856, PWY-5855, and PWY-5857)), heme *b* (HEME-BIOSYNTHESIS-II) and acetaldehyde biosynthesis pathways (PWY66-21 and PWY-6333 with key metabolite: acetyl-CoA and acetaldehyde), L-cysteine degradation pathway (PWY-5329 with key metabolite: pyruvate), NAD phosphorylation pathways (NADPHOS-DEPHOS-PWY and NADPHOS-DEPHOS-PWY-1 with key metabolite: NAD^+^, NADP^+^, and NADPH), and N-end rule pathways (PWY-7801 and PWY-7802 with key metabolite: N-terminal arginyl-protein). Besides, several bacterial lipids metabolic pathways, including oleate and α-Kdo-(2->4)-α-Kdo-(2->6)-lipid IVA (PWY-7664 and PWY-8074 with key metabolite: oleate and α-Kdo-(2->4)-α-Kdo-(2->6)-lipid IV_A_), and some cofactors biosynthesis pathways, e.g., tetrahydromonapterin (PWY0-1433) and molybdenum cofactor biosynthesis (PWY-8171), were also significantly positively associated with *L. marina* development. The isolates belonging to Actinobacteria, Bacteroidetes, and Firmicutes lack these pathways, which might cause *L. marina* developmental delay. In addition, we found that five bacterial metabolic pathways were negatively correlated with *L. marina* development, notably, biosynthesis of pyridoxal 5’-phosphate (PWY-6466), which was reported to be critical for bacterial pathogenicity^32^. The other four negatively correlated metabolic pathways were PWY0-1517, PWY-7247, PWY-6383, and PWY-5532, respectively (Table S3E).

**Fig. 4.**
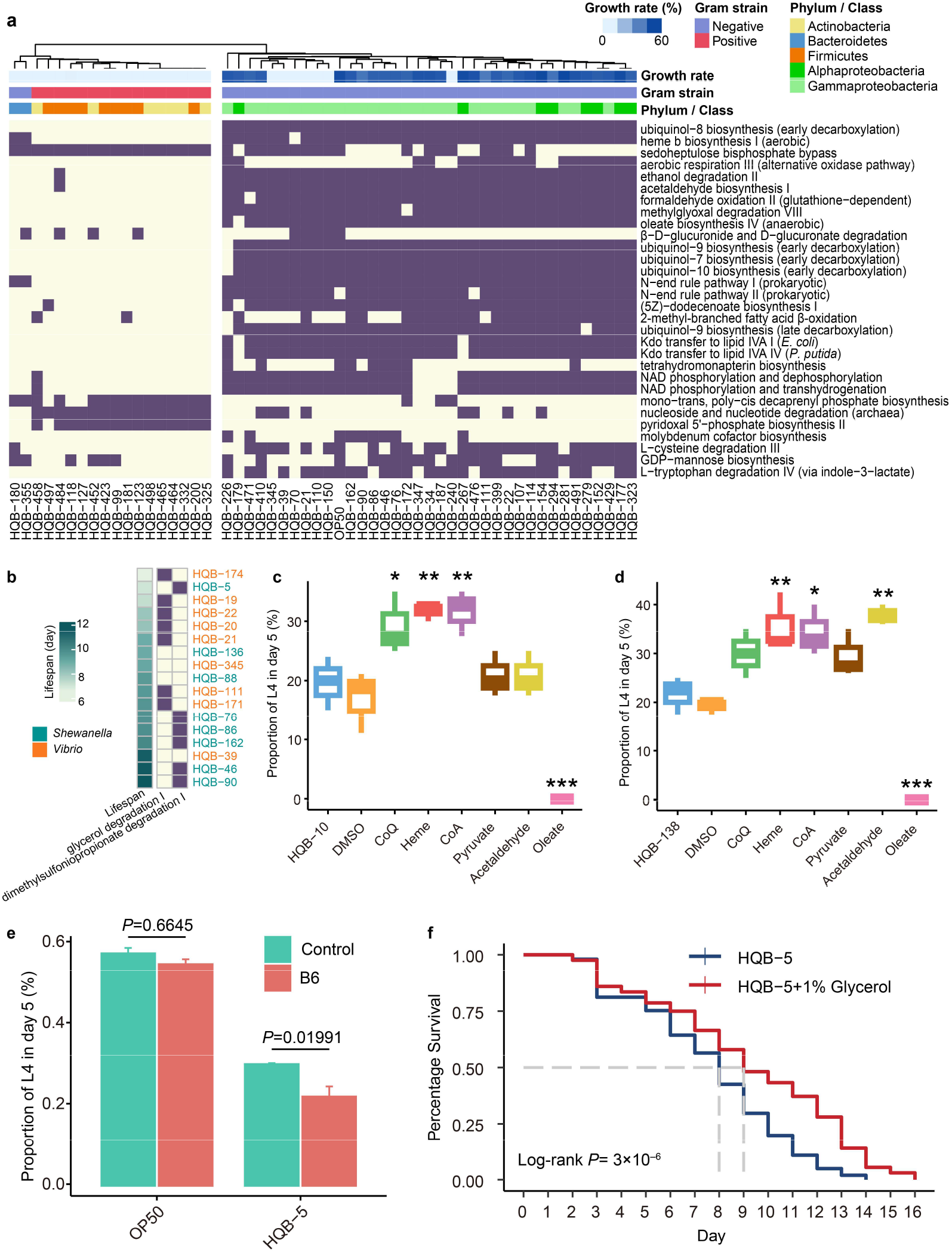
Bacterial metabolites modulate *L. marina* development and longevity. **a**, Bacterial metabolic pathways correlated with *L. marina* development. The developmental rate of *L. marina* (percentage of L4 after 5 days post-hatching), Gram-stain, and phylum/class are indicated by several annotation columns at the top of the matrix. **b**, Bacterial metabolic pathways correlated with *L. marina* lifespan. **c–d,** Metabolites supplementation in HQB-10 (**c**) and HQB-138 (**d**) demonstrates their effects on *L. marina* development. Statistical analyses were performed by one-way analysis of variance (ANOVA) with Dunnett’s multiple comparison test, \**p* < 0.05, \*\**p* < 0.01, and \*\*\**p* < 0.001. Boxes indicate the first and third quartile of counts, lines in boxes indicate median values, and whiskers indicate 1.5 × IQR in either direction. **e,** Vitamin B6 supplementation caused *L. marina* developmental delay. Statistics were performed by Fisher’s exact test. **f,** 1% Glycerol supplementation significantly extended the lifespan of *L. marina* fed *S. algae* HQB-5. Log-rank test was applied for the significance.

To ask if any of the key metabolites identified from the above random forest analysis, could modulate *L. marina* development, we evaluated the single metabolite effect through dietary supplementation. Strikingly, we found that CoQ_10_, heme, and acetyl-CoA supplementation significantly promoted *L. marina* development fed *Alkalihalobacillus algicola* HQB-10. Additionally, we found that dietary supplementation of heme, acetyl-CoA, and acetaldehyde significantly promoted *L. marina* growth fed *Metabacillus halosaccharovorans* HQB-138 (Fig. 4c, d). By contrast, we found that pyruvate significantly promoted *L. marina* development fed *E. coli* OP50 (Fig. S13), while dietary supplementation (e.g., CoQ, heme *b*, acetyl-CoA, acetaldehyde, pyruvate) did not significantly affect *L. marina* development fed with *Paenisporosarcina quisquiliarum* HQB-263 or *Staphylococcus warneri* HQB-123, which belong to “beneficial” and “detrimental” category, respectively. Unexpectedly, we found that oleate supplementation significantly attenuated *L. marina* growth rate or led to death (Fig. 4c,d, and Fig. S13). Moreover, as expected, we found that vitamin B6 supplementation significantly slowed the development of *L. marina* fed with *S. algae* HQB-5 (Fig. 4e).

In addition, we found that *L. marina* lifespan was significantly negatively correlated with glycerol degradation I pathway (PWY-4261), while positively correlated with dimethylsulfoniopropionate degradation I pathway (PWY-6046) (Fig. 4b; Table S3K; Fig. S14). Notably, as expected, we found that 1% glycerol supplementation significantly extended the lifespan of *L. marina* fed with *S. algae* HQB-5 but not OP50 (Fig. 4f, Fig. S15b).

### Bacterial metabolic pathways correlated with *C. elegans* development and longevity

Unexpectedly, we found that the effects of marine native bacteria on the development of *L. marina* and its terrestrial relative *C. elegans* were significantly positively correlated (Supplementary Results S1; Fig. S16). Through random forest regression analysis, we identified bacterial metabolic pathways including GDP-mannose biosynthesis (PWY-5659), aerobic respiration III (PWY-4302), heme *b* biosynthesis I (HEME-BIOSYNTHESIS-II), and L-cysteine degradation III (PWY-5329) were significantly correlated with *C. elegans* development (Fig. 5a; Table S3G), which were the only four pathways that also significantly correlated with *L. marina* growth (Fig. 4a; Table S3E,F). Notably, most bacterial metabolic pathways correlated with animal development were largely different between *L. marina* and *C. elegans* (Fig. 4a; Fig. 5a). Especially, we found that several bacterial metabolic pathways were significantly positively correlated with *C. elegans* development, including GDP-mannose biosynthesis (PWY-5659 with key metabolite: GDP-α-D-mannose), superoxide radicals degradation (DETOX1-PWY), three amino acid degradation pathways (β-alanine: PWY-1781; L-alanine: ALADEG-PWY; and L-cysteine: PWY-5329 with key metabolites: pyruvate and acetyl-CoA), aerobic respiration (PWY-4302 with key metabolite: ubiquinol), heme *b* biosynthesis pathways (HEME-BIOSYNTHESIS-II and HEMESYN2-PWY with key metabolites: heme *b*), and branched-chain fatty acid biosynthesis pathways (PWY-8173, PWY-8174, and PWY-8175 with key metabolites) (Fig. 5a; Table S3G). In contrast, UDP-N-acetyl-D-galactosamine biosynthesis II, N-acetylglucosamine degradation I, and mono-trans, poly-cis decaprenyl phosphate biosynthesis pathways were negatively correlated with *C. elegans* egg-laying time (Table S3G). Although the above key metabolites correlated with *C. elegans* development were distributed to different pathways compared with *L. marina* (Fig. 4a; Table S3E), certain metabolites such as CoQ, acetyl-CoA, pyruvate were shared between both nematodes to promote development (Table S3G). Through single metabolite dietary supplementation, we found that supplementation with acetyl-CoA significantly promoted *C. elegans* development fed *Alkalihalobacillus hwajinpoensis* HQB-118, *Staphylococcus warneri* HQB-123, and *Paenisporosarcina quisquiliarum* HQB-263 (Fig. 5c–e), and pyruvate significantly promoted *C. elegans* development fed with HQB-118 (Fig. 5c). Additionally, we found that *C. elegans* lifespan was negatively correlated with adenine salvage (PWY-6610) (Fig. 5b; Table S3L).

**Fig. 5.**
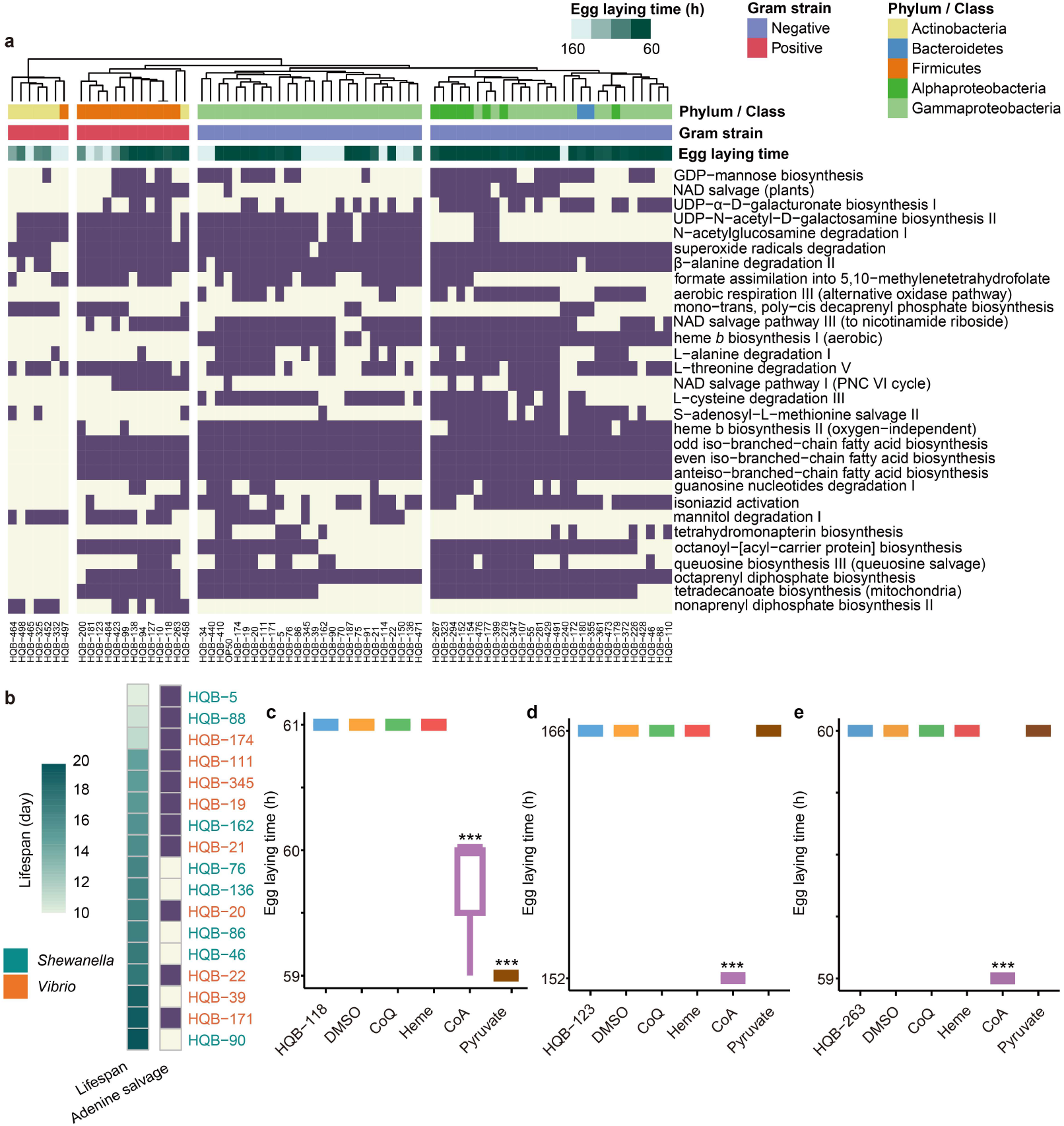
Bacterial metabolic pathways correlated with *C. elegans* developmental rate and longevity. **a,** Bacterial metabolic pathways correlated with *C. elegans* egg-laying time. The pathways were sorted by the importance of random forest regression analysis. Egg-laying time of *C. elegans*, Gram-stain, and phylum/class are indicated by several annotation columns at the top of the matrix. **b,** Bacterial metabolic pathway correlated with *C. elegans* lifespan. The presence of metabolic competence (dark purple in the right panel) was associated with the mean lifespan of *L. marina* (cyan color in the left panel; darker colors indicated increased mean lifespan). Different bacterial genera are indicated by the different colors of the strain names. **c–e,** Metabolite supplementation in strains HQB-118 (**c**), HQB-123 (**d**) and HQB-263 (**e**) demonstrates their effects on *C. elegans* egg-laying time, which is proxies of developmental rates. Statistical analyses were performed by one-way analysis of variance (ANOVA) with Dunnett’s multiple comparison test, \*\*\**p* < 0.001. Boxes indicate the first and third quartile of counts, lines in boxes indicate median values, and whiskers indicate 1.5 × IQR in either direction.

### Gut bacterial colonization and bacterial impacts on *L. marina* and *C. elegans* physiology

To characterize the gut microbiota and its potential effects on animal development and lifespan, we generated germ-free *L. marina* inbred line worms by bleaching eggs to obtain synchronized L1 larvae, which were then fed with the HQbiome bacterial mixture (proportional mixture of each 73 strains; Fig. S18a; Table S4A) on SW-NGM plates. We found that stable *L. marina* gut colonization was observed on day 3 of adulthood (160 h post L1) (Fig. 6a), and the gut microbiome was dominated by *Shewanella* (especially *S. algae*) and *Vibrio* on day 3 and day 5 of adulthood, resembling the surrounding bacterial lawn (Fig. 6a; Table S4B).

**Fig. 6.**
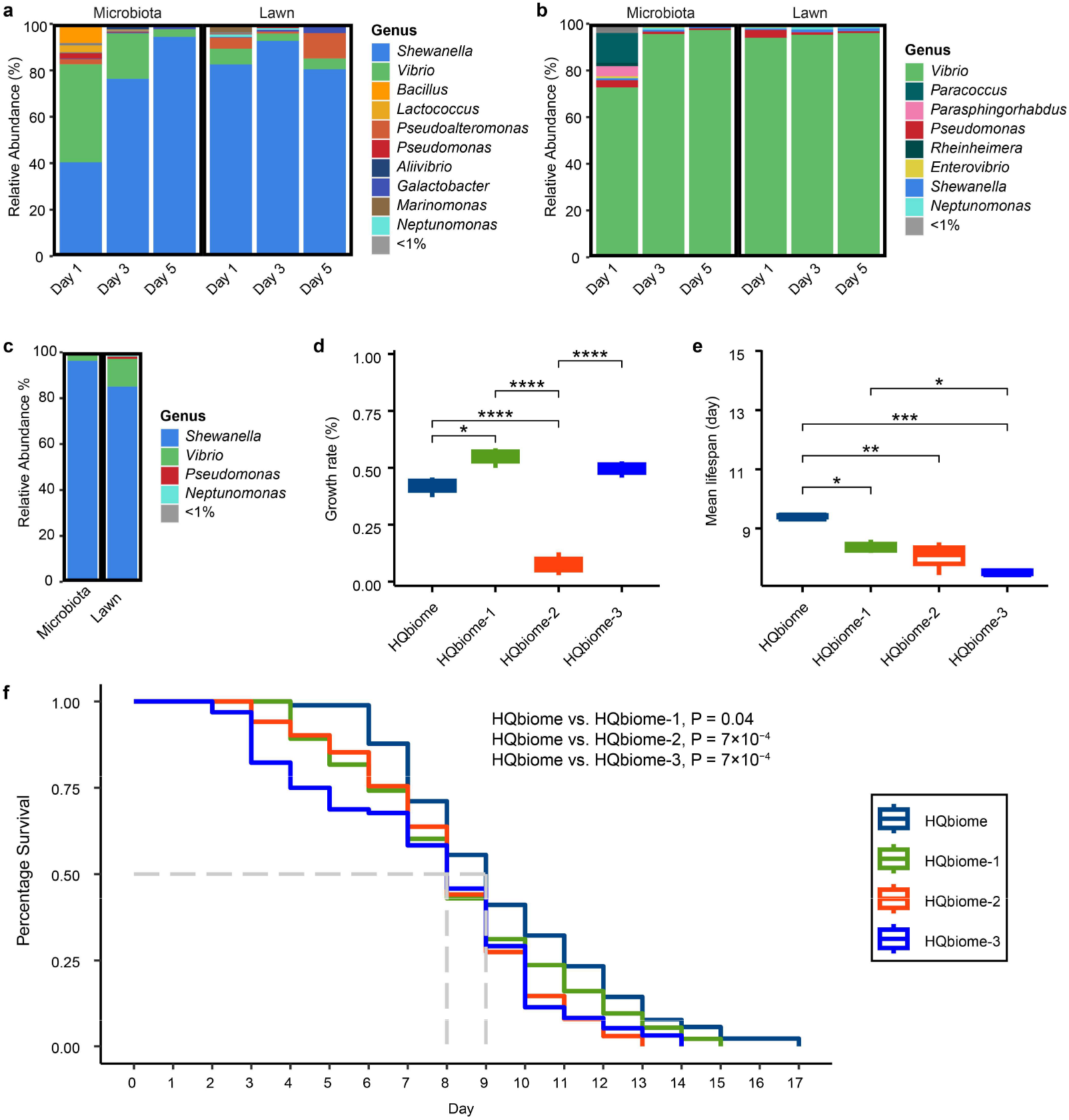
Gut microbiota composition of *L. marina* and bacterial effects on host physiology. **a**, Gut microbiota composition of *L. marina* fed HQbiome. **b**, Gut microbiota composition of *L. marina* fed HQbiome-1. **c**, Gut microbiota composition of *L. marina* fed HQbiome-3 in day 5 of adulthood. **d**, *L. marina* growth rate fed HQbiome, HQbiome-1, HQbiome-2, and HQbiome-3, respectively. **e,** *L. marina* mean lifespan fed HQbiome, HQbiome-1, HQbiome-2, and HQbiome-3, respectively. *P* values in **d** and **e** were generated from one-way ANOVA, followed by Tukey’s post hoc test. \**P* < 0.05; \*\**P* < 0.01; \*\*\**P* < 0.001; \*\*\*\**P* < 0.0001. Boxes indicate the first and third quartile of counts, lines in boxes indicate median values, and whiskers indicate 1.5 × IQR in either direction. **f**, *L. marina* survival curves fed HQbiome, HQbiome-1, HQbiome-2, and HQbiome-3, respectively. Log-rank test was applied for the assessment of significance.

We next asked whether the mixture of the “beneficial” subset HQbiome-1 (Fig. S18b; Table S4A) had similar colonization characteristics compared to the HQbiome mixture. We characterized the gut microbiota of *L. marina* fed HQbiome-1 mixture (proportional mixture of 29 “beneficial” of 73 strains from HQbiome) and found that the gut microbiota stabilization time was similar to that of HQbiome (day 3 of adulthood; Fig. 6a,b). Although there were five *Shewanella* spp. in HQbiome-1 (without *S. algae*), strikingly, we found that *L. marina* gut microbiota was dominated by *Vibrio* (especially *V. parahaemolyticus*) fed HQbiome-1, resembling the bacterial lawn (Fig. 6b; Table S4C). Given that the gut microbiota was dominated by *S. algae* when *L. marina* was fed with HQbiome, and *S. algae* is not available in HQbiome-1, to ask whether *S. algae* play a major role in gut microbiota colonization, we added *S. algae* HQB-5 to HQbiome-1 mixture (termed HQbiome-3, Fig. S18c) and characterized the bacterial composition of lawn and gut microbiota composition (Fig. 6c). Of note, the gut microbiota reverted to be dominated by *Shewanella* when *L. marina* was fed with HQbiome-3 (Fig. 6c), compared with being dominated by *Vibrio* when fed with HQbiome-1 (Fig. 6b). These data suggested that *S. algae* played a major role in *L. marina* gut microbiota colonization.

To ask if the gut microbiota modulates the physiology of *L. marina*, we examined the developmental rate and lifespan when *L. marina* was grown in HQbiome (73 isolates), HQbiome-1 (29 “beneficial” isolates), HQbime-2 (19 “detrimental” isolates), and HQbiome-3, respectively. We observed that the developmental rate of *L. marina* fed HQbiome-1 was significantly faster than that of HQbiome and HQbiome-2, and HQbiome significantly promoted *L. marina* development compared to HQbiome-2, while the developmental rate was similar when fed HQbiome-1 and HQbiome-3 (Fig. 6d). Interestingly, we found that the lifespan of *L. marina* fed HQbiome was significantly longer than that of HQbiome-1 and HQbiome-2, while HQbiome-1 and HQbiome-2 led to similar lifespan (Fig. 6e,f). Of note, we found that HQbiome-3 significantly shortened *L. marina* mean lifespan compared to HQbiome-1 feeding (Fig. 6e), indicating that *S. algae* played a critical role in *L. marina* longevity regulation.

To test whether there are similar gut microbiota characterization and bacterial effects on the development and lifespan between *L. marina* and its terrestrial relative *C. elegans*, we grew *C. elegans* on HQbiome, HQbiome-1, HQbiome-2, and HQbiome-3, respectively. Similar to *L. marina*, we observed that *C. elegans* gut colonization could be stabilized on day 5 of adulthood (155 h post L1) fed either with HQbiome or HQbiome-1 (Fig. 7a,b). In contrast to the observation that *L. marina* gut microbiota was similar in composition to their bacterial lawn (Fig. 6a–c), we found that *C. elegans* gut microbiome composition exhibited selectivity from their bacterial lawn (Fig. 7a–c; Fig. S19). Especially, the gut microbiota of *C. elegans* grown on HQbiome was dominated by *Shewanella* and *Pseudomonas* on day 5 and day 7 of adulthood (Fig. 7a), while *C. elegans* fed with HQbiome-1 was dominated by *Pseudomonas* and *Vibrio* (Fig. 7b). Notably, we observed that *C. elegans* gut microbiota was dominated by *Shewanella* and *Pseudomonas* when animals were grown in HQbiome-3 (Fig. 7c), indicating that *S. algae* played a critical role in *C. elegans* gut microbiota colonization. Similar to *L. marina*, we observed that HQbiome-1 significantly shortened *C. elegans* egg-laying time compared to HQbiome and HQbiome-2, whereas HQbiome-2 significantly delayed egg-laying time compared to HQbiome (Fig. 7d). Of note, *C. elegans* exhibited significant shorter egg-laying time fed HQbiome-3 compared to HQbiome-1, suggesting that *S. algae* played an essential role in promoting *C. elegans* development. By contrast, we found that the lifespan of *C. elegans* fed HQbiome-2 was significantly increased than HQbiome, HQbiome-1, and HQbiome-3, indicating that tradeoffs occurred between *C. elegans* developmental rate and lifespan (Fig. 7d–f). Collectively, these data showed that the impacts of three bacterial mixtures (HQbiome, HQbiome-1, and HQbiome-3) on *C. elegans* developmental rate were in similar trends to that of marine nematode *L. marina*, whereas the lifespan of nematodes might be not only determined by the dominant gut bacteria but also by the interactions between microbiota members in a complex gut bacterial ecosystem.

**Fig. 7.**
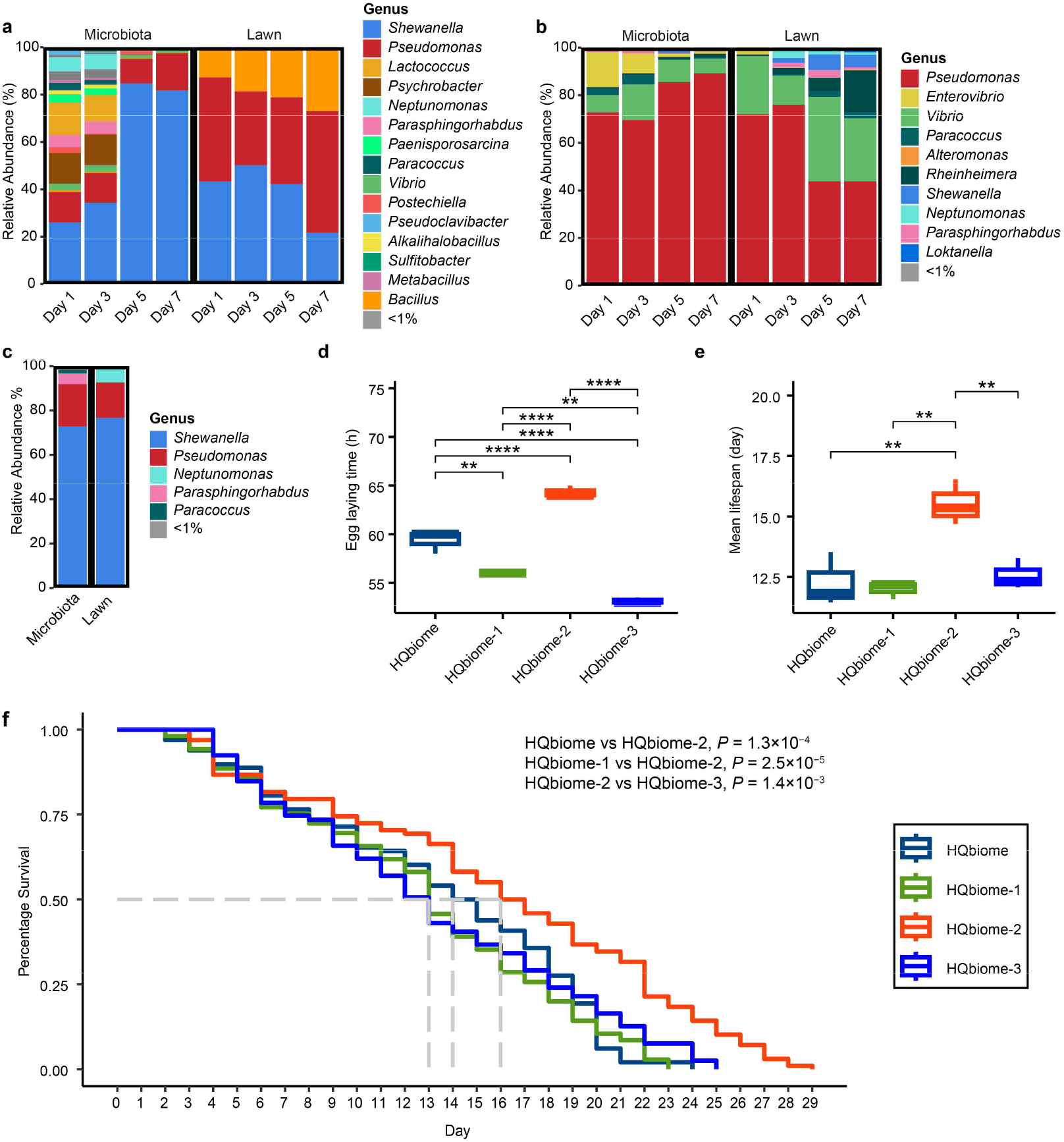
Gut microbiota composition of *C. elegans* and bacterial effects on host physiology. **a**, Gut microbiota composition of *C. elegans* fed HQbiome. **b**, Gut microbiota composition of *C. elegans* fed HQbiome-1. **c**, Gut microbiota composition of *C. elegans* fed HQbiome-3 in day 5 of adulthood. **d**, *C. elegans* egg-laying time fed HQbiome, HQbiome-1, HQbiome-2, and HQbiome-3, respectively. **e,** *C. elegans* mean lifespan fed HQbiome, HQbiome-1, HQbiome-2, and HQbiome-3, respectively. *P* values in **d** and **e** were generated from one-way ANOVA, followed by Tukey’s post hoc test. \**P* < 0.05; \*\**P* < 0.01; \*\*\**P* < 0.001; \*\*\*\**P* < 0.0001. Boxes indicate the first and third quartile of counts, lines in boxes indicate median values, and whiskers indicate 1.5 × IQR in either direction. **f,** *C. elegans* survival curves fed HQbiome, HQbiome-1, HQbiome-2, and HQbiome-3, respectively. Log-rank test was used for the assessment of significance.

## Discussion

In the present study, we applied both cultivation-independent and cultivation-based approaches to characterize the natural sedimentary bacterial composition in Huiquan Bay, where marine nematode *L. marina* was collected. We demonstrated that four phyla were dominant, including Proteobacteria, Bacteroidetes, Cyanobacteria, and Actinobacteria, in accordance with the prominent habitat microbes of terrestrial nematode *C. elegans* (Proteobacteria, Bacteroidetes, Firmicutes, and Actinobacteria)^15^. At the family level, Flavobacteriaceae was abundant in *L. marina* native habitats (8.86%), similar to that in *C. elegans* habitats (3.16%)^15^. Among the total OTUs from the cultivation-independent full-length 16S rRNA gene sequencing, only 0.95% were recovered in our cultivation-dependent bacterial isolation, which was coincident with earlier studies that <1% of marine environmental bacteria were culturable^33^. Notably, through the cultivation-dependent approach, we isolated 539 natural habitat bacteria, including 28 new species.

Both marine nematode *L. marina* and its terrestrial relative *C. elegans* belong to the same family, Rhabditidae^26^. The comparative investigation of *L. marina* and *C. elegans* could provide valuable insights into the adaptations of nematodes to their natural bacterial environments^27^. We examined the effects of 448 marine native bacteria isolates, which were taxonomically similar to the habitat microbiome of *C. elegans* at the phylum level, on the development of *C. elegans*. We found that approximately 82% of single isolates were able to support the *C. elegans* growth, while 18% of isolates significantly delayed *C. elegans* development (Fig. S20). By contrast, we observed that approximately 52.34% of 448 native bacteria supported *L. marina* development, while 47.66% caused *L. marina* developmental delay or death (Fig. 2a). Consistent with a previous report that about 80% of natural habitat-bacterial isolates support *C. elegans* growth, our data indicated that *C. elegans* could survive in a wider range of bacterial community niches. Although a higher percentage of bacteria isolates support *C. elegans* growth, when comparing all 448 individual bacteria effects on *L. marina* developmental rate and the egg-laying time of *C. elegans*, we found that they were positively correlated (Fig. S16; Supplementary Results 1), indicating the conservational role of marine native bacteria in both nematode’ physiology.

Among the 73 HQbiome isolates, 66% (40% “beneficial” and 26% “intermediate”) supported *L. marina* development, while 34% of the isolates were detrimental to *L. marina* development (Fig. 2b). Similar to the effects of 448 marine native bacteria on *C. elegans* development, 77% of HQbiome isolates (68% “beneficial” and 9% “intermediate”) supported *C. elegans* growth, while 23% of the isolates slowed *C. elegans* development. In line with 448 isolates’ effects on *L. marina* growth rate and the egg-laying time of *C. elegans*, HQbiome isolates affect the development of both nematodes in a similar way (Fig. S21).

Based on our reconstructed metabolic models from WGS of 73 HQbiome isolates and their effects on the development of *L. marina*, we found that several metabolic pathways were significantly positively correlated with *L. marina* development rate (Fig. 4a). Essential metabolites from the above pathways included CoQ, heme *b*, acetyl-CoA, acetaldehyde, pyruvate, and oleate. In contrast, we found that the siderophores, pyridoxal 5’-phosphate, and terpenes were negatively correlated with animal growth (Fig. 4a, Fig. S22d, and Fig. S23c). Notably, CoQ_10_, heme, and acetyl-CoA supplementation significantly promoted *L. marina* development fed *A. algicola* HQB-10, and dietary supplementation of heme, CoA, and acetaldehyde significantly promoted *L. marina* growth fed *M. halosaccharovorans* HQB-138 (Fig. 4c,d), in accordance with lack of the corresponding metabolic pathways in both bacteria. Unexpectedly, we found that oleate supplementation caused *L. marina* developmental delay or death (Fig. 4c, d, and Fig. S13). CoQ plays an essential role in mitochondrial energy metabolism^34^ and antioxidant activity^35,36^. Mutations in *C. elegans clk-1* led to synthetic CoQ defect, showing development delay, reduced respiration, and increased life span^37–40^. Heme is a readily bioavailable source of iron for animal life^41^. Heme *b* was reported to be present in the respiratory complexes to facilitate the movement of electrons through the electron transport chain (ETC), and to stabilize ubisemiquinone radicals and decrease the production of reactive oxygen species (ROS)^42^. Given heme could not be synthesized by many nematodes, such as *C. elegans*, which requires environmental heme for growth and development^41^, our data indicated that *L. marina* could not synthesize heme *b* as well. The production of vitamin B6 was reported to be essential for bacterial pathogenicity to *C. elegans*^32^, which might delay *L. marina* development due to increased susceptibility to pathogen bacteria. As expected, pyridoxal vitamin B6 supplementation significantly attenuated the development of *L. marina* fed with *S. algae* HQB-5 (Fig. 4e), which lacks pyridoxal 5’-phosphate biosynthesis pathways.

Only four bacterial metabolic pathways, including GDP-mannose biosynthesis, aerobic respiration III, heme *b* biosynthesis I, and L-cysteine degradation III, were significantly positively correlated with *L. marina* developmental rate and *C. elegans* egg-laying time. However, most of the correlated metabolic pathways were not shared by both nematodes. In contrast to the pro-development role of heme in *L. marina*, heme dietary supplementation did not affect *C. elegans* egg-laying time. The mechanisms by which single bacterial metabolite regulates nematode development deserved to be further explored.

Our study demonstrated that the glycerol degradation I pathway was significantly negatively correlated with *L. marina* lifespan, it has been reported that the dietary supplement of glycerol extended the lifespan in both yeast and rotifers^43,44^. We speculated that the bacteria that lack glycerol degradation competence may supplement *L. marina* with more glycerol and extend the nematode lifespan. In line with our hypothesis, we observed that dietary supplementation of 1% glycerol significantly promoted the longevity of *L. marina* fed *S. algae* HQB-5 but not OP50 (Fig. 4f). By contrast, 1% glycerol supplementation did not extend *C. elegans* lifespan fed either OP50 or *S. algae* HQB-5 (Fig. S15a,c). Several studies in *C. elegans* have uncovered that glycerol supplementation shortened worm lifespan^45,46^. The underpinning mechanisms by which glycerol supplementation regulates *L. marina* longevity deserved to be further explored.

A recent report showed that *C. elegans* gut microbiota stabilized at day 3 of adulthood^14^, while our data revealed that the gut microbiota stabilization occurred at 160 and 150 h post L1 larvae for *L. marina* and *C. elegans*, respectively (i.e., at day 3 of adulthood for *L. marina* and day 5 of adulthood for *C. elegans*), suggesting that distinct environmental bacterial composition may affect the stabilization time of gut microbiota colonization. It was reported that *C. elegans* exhibited three distinct microbiota selection strategies (strong selection, medium selection, and without selection), and the wild-type N2 worms showed medium selection^14^. In agreement with this report, we found that gut microbiota colonization in *C. elegans* N2 worms exhibited medium selection (Fig. 7a,b, and Fig. S19). It was described that *C. elegans* gut microbiota selection was determined either by insulin receptor gene *daf-2* or the TGFβ/BMP-like ligand gene *dbl-1*^14,23^. However, we found that the gut microbiota of *L. marina* was similar in composition to the bacterial lawn without selection (Fig 6a,b and Fig. S19), implying that certain selection might occur in *daf-2* and *dbl-1* and other host genes in *L. marina* genome^14,23^. The gut microbiota was dominated by *S. algae* when *L. marina* was grown on HQbiome, and *Vibrio* was the dominant microbe when fed on HQbiome-1 where *S. algae* is unavailable. Strikingly, the gut microbiota reverted to be dominated by *Shewanella* spp. when *L. marina* was fed with HQbiome-3 (HQbiome-1 plus *S. algae*). These data suggested that *S. algae* played a major role in *L. marina* gut microbiota colonization. Notably, we found that *S. algae* also played an essential role in *C. elegans* gut microbiota colonization. Further studies to elucidate the casual interactions between habitat microbiome, gut microbiota and host biology will provide novel insights into the conservation and management of ecosystems.

Nematodes have evolved through numerous habitat transitions between ocean and land followed by moderate diversification, it was described that *L. marina* evolved from a transition from terrestrial to marine habitats^47^. Further microbiome-mediated comparative functional analysis between marine nematode *L. marina* and its terrestrial relative *C. elegans*, will permit future microbiota manipulation experiments to decipher how natural habitat microbes shape animal host fitness and their evolutionary transition trajectories.

## Materials and Methods

### Full-length 16S rRNA gene sequencing

Samples were collected from Huiquan Bay, Qingdao, China, from June 2020 to June 2021. Full-length 16S rRNA gene sequencing was applied to 80 sedimentary samples. Details can be found in Supplementary Methods.

### Cultivation-dependent bacterial isolation

Intertidal sediments from Huiquan Bay, Qingdao, China (36.05 N, 120.33 E) were collected every about two months over about three-year period (August 2019 to April 2022; Fig. S1). Details of the isolation, sequencing, and media can be found in Supplementary Methods.

### Nematodes maintenance

*Litoditis marina* was a 23^rd^ generation inbred line generated by consecutive full-sibling crosses in our lab^26^. The *Caenorhabditis elegans* N2 strain utilized in this study was obtained from the *Caenorhabditis* Genetics Center (CGC, https://cgc.umn.edu/). *C. elegans* strains were grown on nematode growth media (NGM), while *L. marina* strains were maintained on seawater nematode growth media (SW-NGM)^26^. For regular maintenance, both NGM and SW-NGM were seeded with *E. coli* OP50 as a food source and worms were cultured at 20 °C.

### Development and longevity assays

To assess the influence of different natural bacterial isolates on the development of *L. marina*, we examined the growth rate of *L. marina* on single and mixed bacterial lawns. To assess the influence of 448 bacteria isolates on *C. elegans* development, we examined the egg-laying time of wild-type *C. elegans* N2. The lifespan assay was performed as previously reported^48^. Details can be found in Supplementary Methods.

### Whole genome sequencing and analyses

We generated whole genome sequences for 72 bacterial isolates from HQbiome (73 natural bacteria), of which, 15 were sequenced using both Oxford Nanopore Technologies and Illumina technologies, one with both Pacific Biosciences and Illumina technologies. The remaining 56 strains were sequenced with Illumina only. Details of whole genome sequencing can be found in Supplementary Methods.

### Metabolic network reconstructions

The whole genome sequences were used to reconstruct genome-scale metabolic models using the gapseq v1.2 analysis pipeline^49^. Details of metabolic network reconstructions, BIOLOG verification can be found in Supplementary Methods.

### Random forest regression analysis

We analyzed the association between phenotypic measurements (i.e., the developmental rate and lifespan) and bacterial metabolic capacities using Spearman rank correlation and random forest regression analysis as previously described^22^. A random forest regression model was performed to select features via a permutation-based score of importance using the R package VSURF v1.1.0^50^ by default settings (ntree = 2,000, mtry = p/3).

### Preparation of microbiome mixtures and Gut microbiome composition

In brief, all HQbiome strains were grown on 2216E or R2A plates and incubated at 28 °C until single colonies were visible. The colonies on the plate were then grown overnight at 28 °C and 220 rpm shaking in 15 mL centrifuge tubes (ExCell Bio, China) filled with 10 mL 2216E/R2A. Cultures were harvested by centrifugation at 4,000 *g* for 10 min, adjusted to an OD_600_ of 0.1 in sterile filtered M9 buffer. Inoculums of bacterial mixtures were prepared by combining equal volumes of each isolate, which was then seeded (15 μL) on NGM and SW-NGM plates. Seeded plates were grown overnight at room temperature before use. Gut microbiome samples were collected following the previously published protocol^14^. Details of sample collection and sequencing can be found in Supplementary Methods.

